# Neuronal plasticity at puberty in hypothalamic neurons controlling fertility in female mice

**DOI:** 10.1101/2024.10.06.616855

**Authors:** Yuanxin Zhang, Leonie M. Pakulat, Elisa Galliano, William H. Colledge, Susan Jones

## Abstract

Puberty is a critical transition period to achieve fertility and reproductive capacity in all mammalian species. At puberty, the hypothalamic-pituitary-gonadal (HPG) is activated by neuroendocrine changes in the brain. Central to this are *Kiss1* neurons that produce kisspeptin, a neuropeptide which is a potent stimulator of gonadotropin releasing hormone (GnRH) secretion. *Kiss1* neurons in the arcuate region of the hypothalamus (*Kiss1*^*ARC*^) increase pulsatile secretion of GnRH at puberty. Other developmental maturational changes in the brain are often accompanied by neuronal plasticity changes but this has not been studied in *Kiss1* neurons. Electrophysiological characterisation of *Kiss1*^*ARC*^ neurons from female mice shows that these neurons undergo profound intrinsic plasticity at puberty with a critical window between 3 and 4 weeks. Immature *Kiss1*^*ARC*^ neurons cannot sustain depolarisation-evoked firing for even 500 ms and instead fire a brief burst of high frequency spikes before falling silent. This would make them unsuitable for the sustained activity that is needed to activate GnRH neurons and trigger LH secretion in the HPG axis. After puberty, sustained firing can be maintained, which endows post-puberty *Kiss1*^*ARC*^ neurons with a mature physiological phenotype that is amenable to neuropeptide modulation for generation of burst firing and pulsatile release of kisspeptin. There is a corresponding decrease in the threshold for action potential initiation, a more hyperpolarised post-spike trough and a larger medium after-hyperpolarisation (mAHP). Gene expression analysis showed a significant decrease in *Scn2a* (Na_v_1.2 channel), *Kcnq2* (K_v_7.2 channel) and *Lrrc55* (BK channel auxiliary γ3-subunit) expression and an increase in *Hcn1* (hyperpolarization activated cyclic nucleotide-gated potassium channel) expression which may contribute to the observed electrophysiological changes. Ovariectomy and β-estradiol replacement defined a window of estrogen-dependent plasticity of action potential firing at puberty, such that post-puberty *Kiss1*^*ARC*^ neurons achieve a mature physiological phenotype for activation of the HPG axis.

## Introduction

Puberty is required for an animal to achieve fertility and reproductive capacity to allow continuation of the species. In females, the end of puberty is marked by the establishment of a regular estrous cycle to allow the periodic release of eggs. The estrous cycle is regulated by hormonal feedback loops within the hypothalamic-pituitary-gonadal (HPG) axis. Gonadal sex steroids regulate the output of gonadotropin releasing hormone (GnRH) from the hypothalamus to control release of the gonadotropic hormones LH (luteinizing hormone) and FSH (follicle stimulating hormone) from the anterior pituitary (1). The gonadotropic hormones act on the gonads to stimulate gametogenesis and the production of sex hormones. The feedback mechanism by which sex steroids regulate GnRH secretion is indirect and mediated by kisspeptin neuropeptides released from *Kiss1* neurons. *Kiss1* neurons are found in two main areas of the hypothalamus, the arcuate nucleus (ARC) and the rostroventral periventricular region of the third ventricle (RP3V) which includes the anteroventral periventricular region (AVPV). Kisspeptins are potent stimulators of GnRH secretion, acting via a specific receptor (KISS1R) expressed by GnRH neurons. Mutations in either the *Kiss1r* or *Kiss1* genes cause hypogonadotropic hypogonadism in humans (2-4) and in rodent models (3, 5, 6) characterised by reduced gonadotropin hormone concentrations, pubertal failure, and infertility. However, there is currently no mechanistic understanding of how *Kiss1* neurons trigger the onset of sexual maturity.

The biophysical properties of *Kiss1*^*ARC*^ neurons in adult (fertile) mice have been well described in *ex vivo* brain slices: they show either no firing, regular firing or burst firing (7-14). Critical ion channel genes underlying the electrical properties of adult *Kiss1*^*ARC*^ neurons include *Hcn1* and *Hcn2* (Hyperpolarization-activated, Cyclic Nucleotide Gated channels), *Cacna1g* (Cav3.1, T-type calcium channel) (15), *Trpc5* (Canonical Transient Receptor Potential 5) and G protein-coupled GIRK channels (16) as well as *Kcnd2* (Kv4.2 potassium channels) (14). In contrast, the firing properties of *Kiss1*^*ARC*^ neurons in immature (infertile) mice have not been well characterised. Despite the crucial role of *Kiss1* neurons in puberty, and reports of a modest increase in the number of *Kiss1* immunoreactive cells in the ARC in ewes (17), mice (18) and rats (19), *Kiss1* neuronal plasticity during this critical period remains largely unexplored. A strengthening of synaptic contacts between Kiss1 and GnRH neurons has been reported during puberty in female mice (20). Given that *Kiss1*^*ARC*^ neurons in adult mice must fire action potentials for 2–5 min at a minimum of 10Hz to increase LH to physiological levels (21), it is of particular interest to understand whether and how *Kiss1*^*ARC*^ neurons change at puberty. Intrinsic neuronal plasticity refers to changes in the ion channels expressed by a neuron (22, 23) including those underlying the action potential. The biophysical basis of action potential firing reflects the complement of voltage- and calcium-sensitive ion channels in the soma and the AIS (axon initial segment) that influence the number, frequency and pattern of repetitive firing (24).

We hypothesised that immature *Kiss1*^*ARC*^ neurons in pre-puberty female mice are unable to achieve sustained firing, and that neuronal plasticity occurs within *Kiss1*^*ARC*^ neurons during puberty to attain a mature intrinsic physiological phenotype. Our results confirm that immature *Kiss1*^*ARC*^ neurons cannot sustain depolarisation-evoked firing for even 500 ms and instead, fire a brief burst of high frequency spikes before falling silent. Maximum spike number increases and maximum spike frequency decreases at puberty, accompanied by a decrease in the threshold for action potential initiation, a more hyperpolarised post-spike trough, a larger medium after-hyperpolarisation, and changes in the expression of several ion channels. The increase in maximum spike number is seen if mice undergo ovariectomy (with estrogen implants) at the end of puberty (6 weeks) but not prior to the start of puberty (3 weeks), suggesting there is an estrogen-dependent window of plasticity between 3-6 weeks.

## Methods

### Animals and brain slice preparation

All animal experimentation was carried out under the UK Home Office Animals (Scientific Procedures) Act 1986, following ethical review by the University of Cambridge. The mice used in this study were derived from a *Kiss1-Cre* colony (25) bred with a tdTomato reporter mouse line (*Gt(ROSA)26Sor–loxSTOPlox– TdTomato*; stock #007909; The Jackson Laboratory) to visualize *Kiss1* neurons. Genotyping was done as described previously (25). Heterozygous *Kiss1-Cre* mice were used which show normal fertility; all mice were homozygous for the tdTomato reporter gene. Mice were group housed by sex (n = 2-5 mice per cage) under a 12:12 h dark/light regime with ad *libitum* access to food and water. Puberty was defined as the initial opening of the vaginal canal (VO). Pre-puberty female mice were separated into two groups: a 3-week-old group (3w: P22-P26) and a 4-week-old group (4w: P28-P34), while post-puberty mice were defined as being 6-8 weeks old (6-8w: P43-P62), with a regular estrous cycle, which was determined cytological examination of vaginal lavages (26).

Mice were killed by cervical dislocation and brains removed and submerged in ice-cold solution containing (mM): 206 sucrose, 2.5 KCl, 1.25 NaH_2_PO_4_, 26 NaHCO_3_, 20 glucose, 5 MgCl_2_, 1 CaCl_2_; pH 7.4 with 95% O_2_/5% CO_2_. Coronal brain slices (250 µm) through the ARC were prepared using a Campden 7000smz Vibrating Microtome (Campden Instruments, UK). Slices were then transferred to an incubation chamber containing (mM): 119 NaCl, 2.5 KCl, 1.25 NaH_2_PO_4_, 26 NaHCO_3_, 20 glucose, 5 MgCl_2_, 2 CaCl_2_ at 30 °C, saturated with 95% O_2_ and 5% CO_2_, and left undisturbed for at least 60 minutes.

### Electrophysiology

Brain slices were transferred to a recording chamber on the fixed stage of an Olympus BX51WI inverted microscope fitted with Differential Interference Contrast and LEDs and filter cubes to visualize florescence at 554 nm wavelength. Brain slices were continuously perfused at 2-3mL/min with fresh oxygenated solution (as above but with 1 MgCl_2_ and 20 glucose) at 30 ± 2 °C (Warner Instruments). Patch pipettes with a tip resistance of 4-6MΩ were filled with intracellular solution containing (mM): 120 K^+^-Gluconate, 9 KCl, 3.5 MgCl_2_, 4 NaCl, 10 HEPES (4-(2-hydroxyethyl)-1-piperazineethanesulfonic acid), 4 Na_2_ATP, 0.4 Na_3_GTP, 0.5 CaCl_2_, 0.3 EGTA (Ethylene glycol-bis(2-aminoethylether)-*N,N,N′,N′*-tetra-acetic acid), pH 7.2-7.3 (using KOH), 270-285 mOsm. Membrane resistance (R_M_) and membrane capacitance (C_M_) (Table 1) were monitored using a 5mV/ 10ms test pulse between each experimental protocol. Series resistance was typically 10-20 MΩ. Recordings were discarded if series resistance exceeded 25 MΩ or if the input resistance was less than 500 MΩ, or if either of these changed by >20%. *Kiss1*^*ARC*^ neurons, identified by the expression of tdTomato, were voltage clamped to -60mV or held in current clamp at the resting membrane potential or at -75 mV (using current injection if necessary) using an Axopatch 200B patch clamp amplifier (Molecular Devices, USA). To measure action potential firing under standard current clamp conditions, depolarising current was injected in 20 pA increments either for 10 ms (to capture the waveform of a single action potential) or for 500 ms (to determine the number, frequency and regularity of action potential trains). Data were low pass filtered (3 kHz) and acquired to Spike 2 software (Version 10; Cambridge Electronic Design, Cambridge, UK) via a Micro1401 at a sampling frequency of 20 kHz for offline analysis using Spike 2 (Version 10). DataView (version 12.2.3; Dr. W. J. Heitler, University of St Andrews) was used to measure parameters during single action potentials: spike threshold (sudden acceleration of dV/dt), spike peak, post-spike trough (as an indicator of the fast after-hyperpolarisation, AHP) and peak-to-trough time (Table 1). Resting membrane potential (RMP) was recorded in current clamp immediately after whole-cell formation in every experiment. RMP, action potential threshold and trough values (Table 1) have been corrected for a liquid junction potential of -18mV.

**Table 1.** Passive and active membrane properties of *Kiss1*^*ARC*^ neurons from 3, 4 and 6-8 week old female mice (mean ± SE)

|  | 3w | 4w | 6-8w |
| --- | --- | --- | --- |
| RMP mV | -62.9 ± 1.3 | -62.4 ± 1.6 | -65.7 ± 1.3 |
| $R_M$ MOhm | 1115 ± 52.5 | 1031 ± 51.3 | 1089 ± 43.4 |
| $C_M$ pF | 35.1 ± 1.3 | 35.8 ± 1.3 | 34.5 ± 0.9 |
| Maximum spike number | 14 ± 2 | 22 ± 1.7 | 24 ± 1.6 |
| Maximum frequency Hz | 111 ± 6 | 106 ± 9 | 75 ± 4 |
| Maximum duration ms | 354 ± 25 | 434 ± 12 | 457 ± 6 |
| CoV of ISI | 0.29 ± 0.04 | 0.2 ± 0.02 | 0.18 ± 0.02 |
| Rheobase pA | 29 ± 2.5 | 32 ± 2.4 | 33 ± 2.4 |
| Maximum current pA | 70 ± 5 | 101 ± 10 | 113 ± 6 |
| Threshold mV | -49 ± 1.2 | -53 ± 1.4 | -54 ± 0.8 |
| Trough mV | -64 ± 3.5 | -84 ± 1.2 | -85 ± 1.1 |
| Peak mV | 19 ± 2.5 | 25 ± 2.7 | 20 ± 2.1 |
| Peak to trough time ms | 3.1 ± 0.2 | 2.2 ± 0.1 | 2.5 ± 0.15 |
| mAHP amplitude mV | 2 ± 0.2 | 3.9 ± 0.4 | 4 ± 0.26 |

### Ovariectomy

Mice were injected with Metacam (5mg/ml) at a dose of 1mg/kg and induced with 5% isoflurane mixed with oxygen for inhalation to ensure they were pain-free and unconscious. Maintenance anaesthesia consisted of 2-3% isoflurane mixed with oxygen for inhalation. Laparotomy was performed to remove the ovaries from both sides, and the incision sutured and clamped. After the experiment, each mouse received 0.1 ml of Metacam orally for pain relief for 3 days and weight was monitored for 1 week. Estrogen implants were given after OVX (OVX+E) to reinstate estrogen levels (27). To create the estrogen implant, β-estradiol (Sigma, cat# E8875-250MG) was mixed with medical adhesive glue (Dow Corning Silastic Medical adhesive, silicon type A) to a final concentration of 4 µg/ml. This mixture was placed in silastic tubing (ID 1.02 mm, OD 2.16 mm, Corning) and cut into 1 cm lengths after drying in the dark for 24 hours. The estradiol concentration in this 1 cm implant was 4 µg/cm, which should provide sufficient β-estradiol to the OVX mice within the physiological range for a maximum of 1 week. Estradiol levels have been confirmed using these implants to be 28.2 ± 5.6 pg/ml in intact diestrous mice (n = 5) and 33.1 ± 6.5 pg/ml in OVX mice (n = 9) (27). Mice requiring estrogen implants were anesthetized in the same manner as for the OVX surgery. The 1 cm β-estradiol implant was placed into a subcutaneous pocket behind the head, and the incision was closed.

### Cell dissociation and quantitative PCR

Cell dissociation was performed on brain slices containing the arcuate nucleus. Tissue of approximately 2.4 mm diameter was punched out of the arcuate region using Glison P1000 plastic pipette tips. The punches were incubated with papain (Cat No./ID: LK003150, Worthington Biochemical Corporation, 0.6 U) and DNase (15 U) at 37°C for 30 minutes. Following incubation, the punches were washed three times in EBSS#2 (Earle’s Balanced Salt Solution) (containing (2R)-amino-5-phosphonovaleric acid (AP-V), 0.8 mM kynurenic acid, albumin-ovomucoid inhibitor (3.4% (w/v), 3.47 U/ml DNase, and 5% (w/v) trehalose) by gentle pipetting and aspiration of the solution, and finally suspended in a 200 μl drop of the same solution. The tissue was mechanically dissociated by repeated pipetting. To enhance cell viability during manual cell isolation, 300 μl of DMEM/F12 (Phenol Red free) medium with serum (10% (v/v) fetal bovine serum, 10% (v/v) horse serum), 5% (w/v) trehalose, 0.025 mM AP-V, and 0.4 mM kynurenic acid was added to the drop. The cells were transferred to a Petri dish and visualized under a fluorescent microscope (EVOS® FL Auto Imaging System, Cat No./ID: 4471136; ThermoFisher Scientific). Pools of tdTomato positive cells were washed in this buffer to remove unlabelled cells and debris. Around twelve cells were collected as one pool and stored in 10 μl aliquots of DNAse-RNase free H_2_O, along with dithiothreitol (0.1M DTT, ThermoFisher Scientific) and 40 U RNase inhibitor (RiboLock RNase Inhibitor, ThermoFisher Scientific, CAT No. E00381), at -80°C. Cell pools were converted to cDNA using Superscript™ IV Reverse Transcriptase (ThermoFisher, Cat. No. 18090010). Both random hexamers and oligo dT were used as primers. In each conversion from cell pools to cDNA, one pool of cells was processed as a no-RT-cDNA negative control by substituting the Reverse Transcriptase with RNase/DNase-free water.

Quantitative PCR (qPCR) was used to examine the expression levels of selected ion channel genes in *Kiss1*^*ARC*^ neurons. qPCR was conducted using the SSoAdvanced Universal SYBR Green Supermix (Bio-Rad, Cat. No.: 1725271) on a Quant-Studio™ Real-Time PCR System (ThermoFisher, Cat. No.: A28567). The cycling conditions were as follows: an initial denaturation step at 95°C for 2 minutes, followed by 40 cycles of amplification consisting of a denaturation step at 95°C for 5 seconds and an annealing/extension step at 60°C for 30 seconds. Lastly, a melt curve analysis was performed (1 second at 95°C, 20 seconds at 60°C, 1 second at 95°C). The reactions were set up in triplicates, with 1μl of cDNA sample per well. Each plate of reactions included a no-RT control and a negative control (no template), both in triplicate. Quantitation was performed standardised to the house-keeping gene *Tbp* using the ΔΔCt method of Pfaffl (28) and relative to the average Ct values of the pre-puberty samples.

### Statistical analysis

Statistical analyses were performed using GraphPad Prism (version 9.4.1 for Windows, GraphPad Software, San Diego, CA, USA). A Shapiro–Wilk normality test was used to examine if data were normally distributed. Data are presented as mean ± standard deviation (SD) or standard error of the mean (SE). A t-test or Mann–Whitney non-parametric test was used, as appropriate, to compare measures between two groups. A one-way ANOVA followed by a Dunnett’s *post hoc* test or Kruskal-Wallis with Dunn’s *post hoc* test was used to compare the effect of one variable on more than two groups of data. A mixed effects analysis or two-way ANOVA followed by a Sidak’s *post hoc* test was used to compare the effects of two variables; F and P values are given in the Figure Legends. Statistical significance was accepted as p≤0.05. N refers to the number of mice, while n refers to the number of cells (and is equivalent to the number of slices).

## Results

To investigate whether and when intrinsic plasticity happens at puberty in *Kiss1*^*ARC*^ neurons in female mice, whole cell patch clamp recordings were made from these neurons in mice aged 3 week (3w), 4 week (4w), and 6-8 week (6-8w). One possible change might be at the level of the passive properties of the membrane as *Kiss1*^*ARC*^ neurons mature. However, there was no significant difference in RMP, R_M_ or C_M_ in 3w (n=36, N=13), 4w (n=31, N=21) or 6-8w (n=43, N=22) mice (Table 1).

Action potential firing properties were investigated by injecting current to maintain the membrane potential at -75mV (avoiding spontaneous firing) and then injecting incremental current pulses for 500 ms. Depolarising current of 20pA or more was sufficient to evoke at least one action potential in *Kiss1*^*ARC*^ neurons from pre-puberty and post-puberty mice (Figure 1A, B), but the maximum number of action potentials fired significantly changed just prior to puberty onset, at 4w (Figure 1B, C). There was a significant effect of age and current injection on the number of spikes fired (Figure 1B), and the maximum number of spikes that could be evoked was significantly more in *Kiss1*^*ARC*^ neurons from 4w or 6-8w mice compared with 3w mice (Figure 1C; n/N as above). There was a significant effect of age and current injection on the frequency of firing (Figure 1D), with firing frequency decreasing significantly by 6-8w (Figure 1E). The reason for the increase in spike number but decrease in frequency is explained by the increase in the duration of sustained firing during the 500 ms depolarisation (Figure 1A, F) after 4w. Firing pattern, measured as the coefficient of variation in the inter-spike intervals, was significantly less variable in post-puberty mice (Figure 1G). Together, these data suggest that *Kiss1*^*ARC*^ neurons in 3w pre-puberty female mice show brief and irregular high frequency firing, but as they approach puberty their firing patterns stabilise into a lower frequency, regular pattern which they sustain for a longer time.

**Figure 1.**
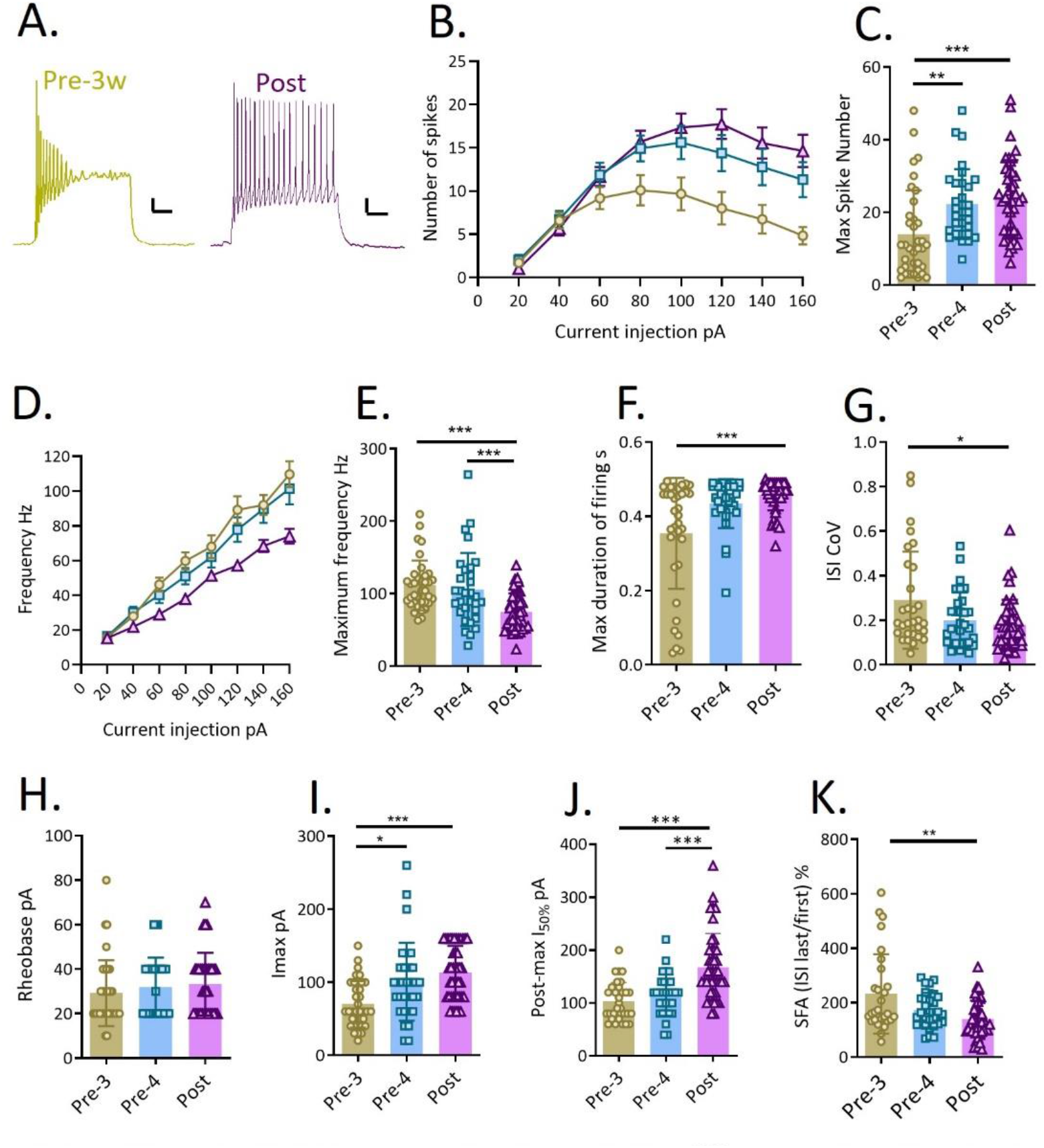
Repetitive action potential firing properties change in *Kiss1*^*ARc*^ neurons from female mice at puberty. A. Example traces of repetitive firing during 500ms current injections in *Kiss1*^*ARc*^ neurons from a 3 week old pre puberty female mouse (left, gold) and a post-puberty female mouse (right, purple). Scale bar: 10mV, 100ms. B. Number of spikes versus increasing current injection. Gold: 3w (n = 36); Blue: 4w (n = 31); Purple: 6-8w (n = 43). Mixed model: Current, P<0.0001, F(7,865)=25; Age, P<0.0001, F(2,850)=27. Tukey’s multiple comparisons show differences between 3w and 4w (P<0.0001) and 3w and 6-8w (P<0.0001). C. Maximum number of spikes, regardless of current injection, from each recording at the three ages as in B. Kruskal-Wallis test, P<0.0001. D. Firing frequency (Hz) versus increasing current injection. Colours and **n** as for B. Mixed model: Current, P<0.0001, F(7,686)=73; Age, P<0.0001, F(l.5,524)=27. Tukey’s multiple comparisons show differences between 3w and 6-8w (P<0.0001) and 4w and 6-8w (P<0.0001). E. Maximum spike frequency, regardless of current injection, from each recording at the three ages as in B. F. Maximum duration of firing during S00ms current injection from each recording at the three ages as B. G. Coefficient of variation in the inter-spike intervals from the maximum number of spikes during each recording at the three ages as in B. H. Rheobase from each recording at the three ages as in B. I. Current required to evoke the maximum number of spikes (up to 160pA) during each recording at the three ages as in B. J. Current required to evoke less than half the maximum number of spikes (post-maximum) during each recording at the three ages as in B. K. Ratio of the last and first inter-spike intervals as an indicator of spike frequency adaptation (SFA) in each recording at the three ages as in B. All *posthoc* P values: *P<0.05; **P<0.01; ***P<0.001.

The amount of current needed to evoke firing (rheobase) during a 500 ms injection did not change at puberty (Figure 1H), although the current inducing the maximum number of spikes (Figure 1I) was significantly larger at 4w and 6-8w, indicating a steadier linear increase in the input-output function in the older mice. Consistent with this, current injections beyond 120 pA led to a progressive decrease in spike number in *Kiss1*^*ARC*^ neurons of all ages, but this was more pronounced in 3w mice, reflected by a larger current needed to reduce the post-maximum spike number by more than 50% at 6-8w (Figure 1J). The ratio of the last: first inter-spike interval in the spike train was significantly lower in *Kiss1*^*ARC*^ neurons from post-puberty compared with pre-puberty mice (Figure 1K), indicating less spike frequency adaptation (SFA) is occurring. Together, these data suggest that *Kiss1*^*ARC*^ neurons from 3w female mice may be unable to fire sustained trains of action potentials due to depolarisation block, spike frequency adaptation, or both.

Changes in the firing properties of *Kiss1*^*ARC*^ neurons at puberty most likely reflect changes in the expression of ion channels underlying neuronal excitability, in particular voltage-gated Na^+^ and K^+^ ion channels (29), and this can be detected in the action potential waveform (24). Incremental current pulses were injected for 10ms to evoke a single action potential (Figure 2A). Waveform parameters such as the spike amplitude, width and fast after-hyperpolarisation (AHP), can be measured relative to the threshold to allow comparisons between the groups. However, there was a significant shift to a more hyperpolarised threshold for spike initiation in 6-8w (n=34, N=18) compared with 3w (n=30, N=12) mice (Figure 2B). Threshold values showed a significant negative correlation with max spike number (Figure 2C). As threshold changed at puberty, the absolute values of the peak, the time from the peak to the trough, and the absolute trough were measured as indicators of spike height, spike repolarisation rate and AHP size respectively. Spike peak was not significantly different (Table 1), but peak to trough time was significantly longer at 3w (n=30, N=12) compared with 4w (n=27, N=18) mice (Figure 2C), and the absolute trough was significantly more hyperpolarised in *Kiss1*^*ARC*^ neurons from 4w (n=26, N=17) and 6-8w (n=18, N=10) mice compared with 3w (n=30, N=12) mice (Figure 2D). Interestingly, there was considerable variation in the trough in the 3w mice. Trough showed a significant negative correlation with maximum spike number (Figure 2E). Thus, a more hyperpolarised threshold and trough were both associated with a higher maximum spike number.

**Figure 2.**
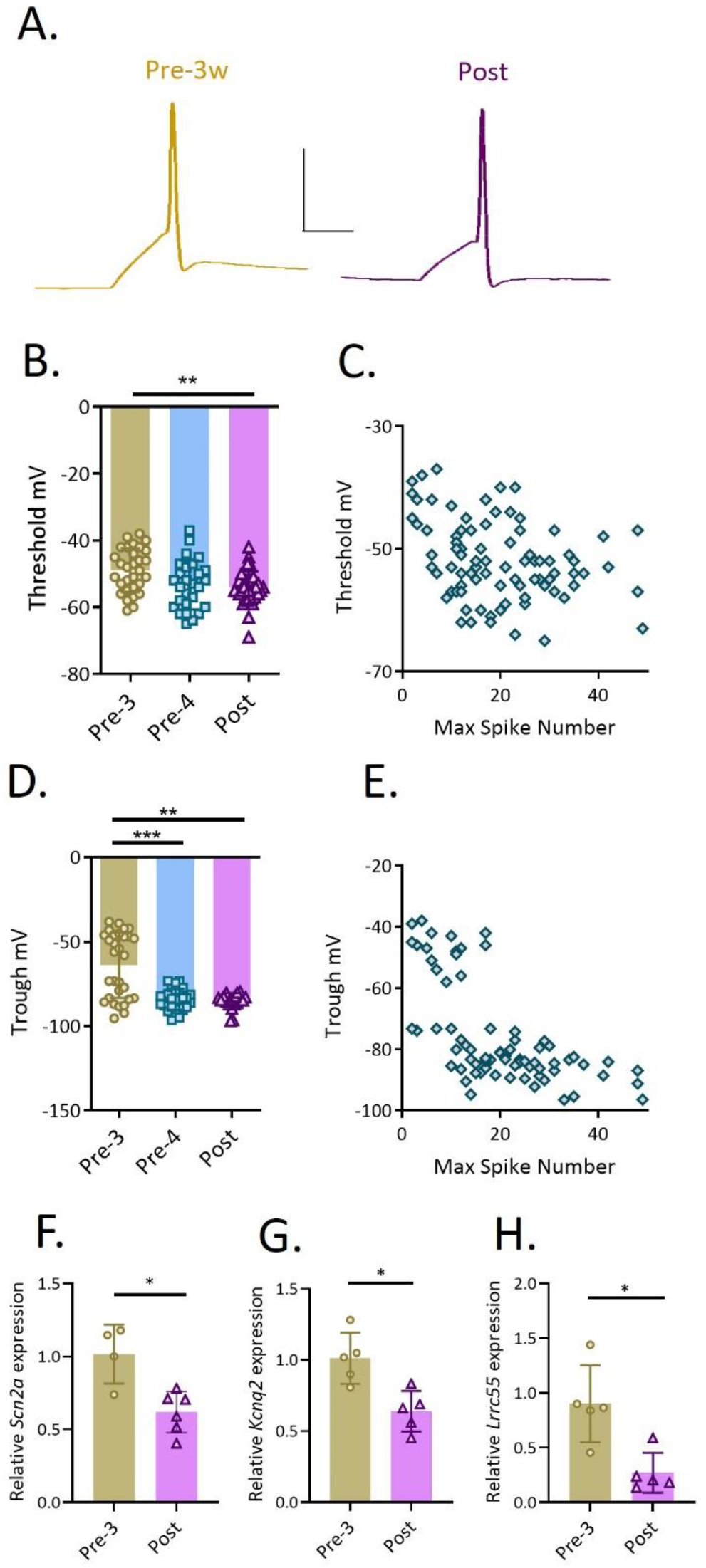
Changes in action potential waveform and ion channel expression in *Kissl*^*ARc*^ neurons from female mice at puberty might contribute to intrinsic plasticity. A. Example traces of single action potentials during 10ms current injections in *Kiss1*^*ARc*^ neurons from a 3 week old pre-puberty female mouse (left, gold) and a post-puberty female mouse (right, purple). Scale bar: 10mV, 10ms. B. The threshold for action potential initiation (mV) at 3w (gold, n = 30), 4w (blue, n = 27) and 6-8w (purple, n = 34). Kruskal Wallis test, P<0.006. C. Significant negative correlation between threshold (mV) and maximum spike number (Spearman r = -0.32; P=0.003). D. Absolute trough value following an action potential (ages and colours as in B; n = 30 (3w), 26 (4w) and 18 (6-8w)). Kruskal-Wallis test, P=0.0001. D. Significant negative correlation between trough (mV) and maximum spike number (Spearman r = -0.66; P<0.0001). F-H. Relative expression of genes encoding Na_v_ l.2 *(5cn2a)*, K_v_7.2 *(Kcnq2)* and the y3 subunit of the BK channel *(Lrrc55)* in isolated *Kiss1*^*ARC*^neurons from 3w pre-puberty and 6-8w post-puberty female mice (Mann-Whitney U test in all cases). All *posthoc* P values: *P<0.05; **P<0.01; ***P<0.001.

Quantitative PCR (qPCR) was used to explore gene expression changes at puberty that could account for the electrophysiological changes, namely those encoding voltage gated sodium (Na_V_) and potassium (K_V_) channels and the large conductance voltage- and calcium-sensitive (BK) potassium channels. Analysis of several house-keeping genes identified the TATA box binding protein gene *Tbp*, as the most appropriate for this comparison as it did not change expression significantly in the *Kiss1*^*ARC*^ neurons at puberty (P = 0.8454).

The expression level of the gene encoding the Na_V_1.2 channel, *Scn2a*, was significantly higher in pre-pubertal *Kiss1*^*ARC*^ neurons (N=4) compared to post-pubertal *Kiss1*^*ARC*^ neurons (N=6; Figure 2F). The expression level of *Kcnq2* (K_V_7.2), which encodes the voltage-gated potassium channel subfamily Q member 2, was significantly higher in pre-pubertal *Kiss1*^ARC^ neurons (N=5), while *Kcnq5* (K_V_7.5) did not show any significant difference (N=4-5; Figure 2G). BK channels can contribute to fast AHPs; the BK*α* subunit (gene: *Kcnma1)*, did not show any significant difference at puberty (P=0.69, N=4), but expression of the BK*γ*3 subunit gene (*Lrrc55*) was significantly higher in pre-pubertal *Kiss1*^*ARC*^ neurons (N=5; Figure 2H).

In addition to the fast AHP occurring immediately after the action potential spike, as shown by the significantly lower trough in Figure 2D, an AHP lasting <300ms was observed following a 500 ms train of action potentials (Figure 3A), referred to as a medium AHP (mAHP) (30). Significantly more post-puberty *Kiss1*^*ARC*^ neurons had a mAHP (Figure 3B). In neurons with a detectable mAHP, the amplitude was significantly larger in post-puberty neurons (Figure 3C). Across all age groups, mAHP amplitude showed a significant negative correlation with firing frequency (Figure 3D): the larger the mAHP, the lower the maximum spike frequency. This suggests that the longer-lasting mAHP could contribute to lower frequency firing in post-puberty *Kiss1*^*ARC*^ neurons. Different ion channels are proposed to contribute to a mAHP, including small conductance calcium activated K^+^ channels (SK), voltage-gated potassium channel subfamily Q (which did not increase at puberty; Figure 2G), and the hyperpolarisation-activated non-selective cation channels, HCN, which contribute to the post-trough depolarisation. The most common SK channels in the brain are SK2 and SK3, whose genes *Kcnn2* and *Kcnn3* showed no significant difference between 3w and 6-8w *Kiss1*^ARC^ neurons (both N=4; Figure 3E, F). The expression of four *HCN* genes was also measured; only the gene that encodes the HCN1 channel was significantly higher in post-pubertal *Kiss1*^ARC^ neurons (N=5; Figure 3G); an increase in post-pubertal *Hcn1* in *Kiss1*^ARC^ neurons might contribute to the recovery from the mAHP. However, the ion channel responsible for the hyperpolarisation during the mAHP remains unknown.

**Figure 3.**
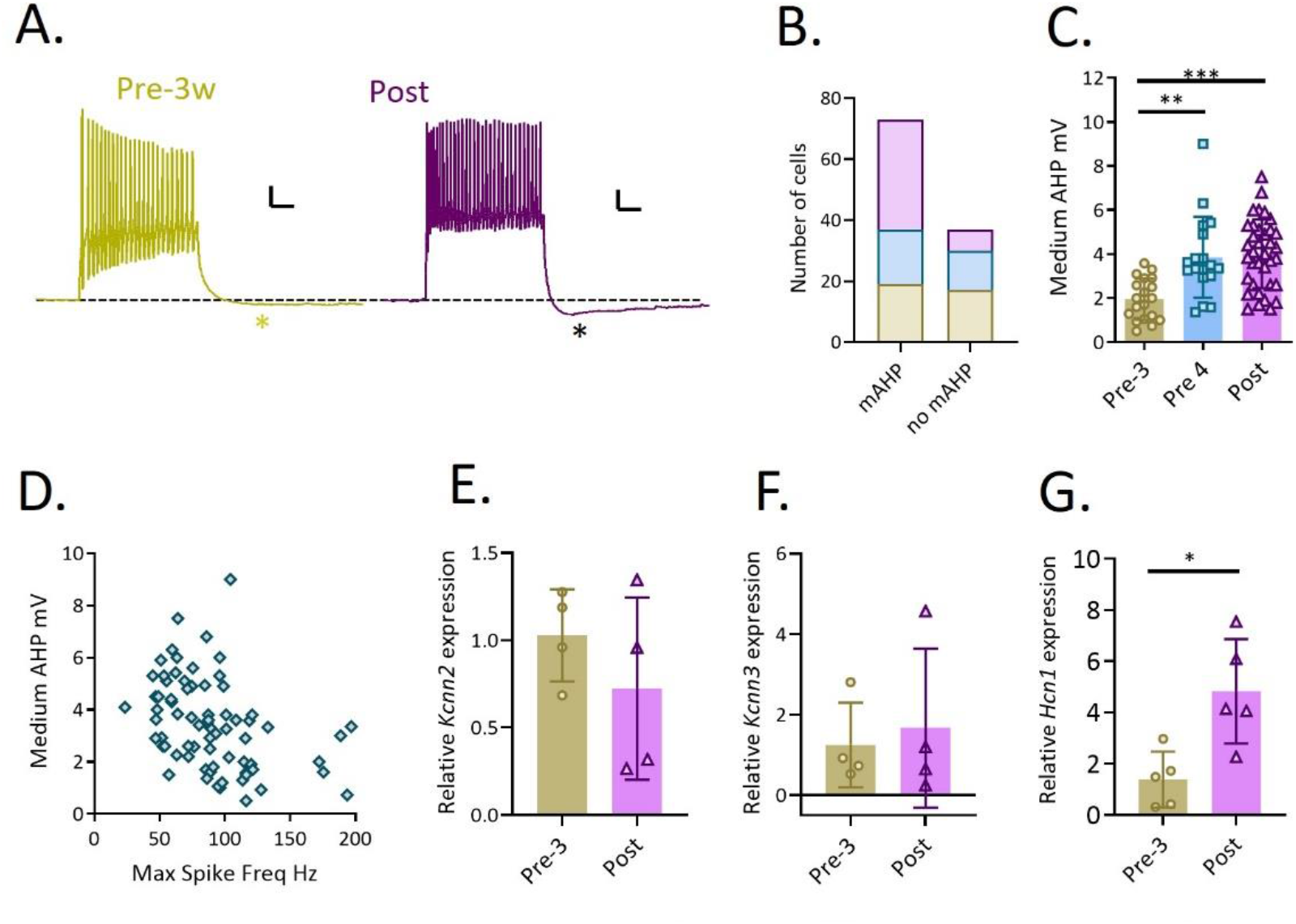
Larger mAHP following a train of action potentials in *Kissl*^ARC^ neurons from post-puberty female mice. A. Example traces of the mAHP (*) following a train of action potentials during 500ms current injections in *Kissl*^*ARC*^ neurons from a 3 week old pre-puberty female mouse (left, gold) and a post-puberty female mouse (right, purple). Scale bar: 10mV, 100ms. B. The number of *Kissl*^*ARC*^ neurons that had a detectable mAHP in 3w (gold, n = 19/36), 4w (blue, n = 18/31) and 6-8w (purple, n = 36/43); Chi Squared test, P=0.008. C. Amplitude of the mAHP (Kruskal Wallis test, P<0.0001). D. Significant negative correlation between mAHP (mV) and maximum firing frequency (Hz) (Spearman r = -0.37; P=0.001). E-G. Relative expression of genes encoding SK2 (*Kcnn2*), SK3 (*Kcnn3*) and the HCN1 (*Hcnl*) in isolated *Kissl*^*ARC*^ neurons from 3w pre-puberty and 6-8w post-puberty female mice (Mann-Whitney U test in all cases). All posthoc P values: *P<0.05; **P<0.01; ***P<0.001.

To investigate the trigger for the change in firing properties at puberty, the role of the sex steroid hormone, estrogen was considered. Estrogen levels begin to rise prior to puberty onset and fluctuate during the estrous cycle after puberty with lowest levels just after ovulation. If increasing levels of estrogen are required for the changes in spike firing after puberty, then differences in firing properties might be expected at diestrous versus estrous, which have different levels of circulating estrogen (31). However, there was no significant difference between estrous and diestrous in maximum spike number or frequency of firing in response to 500 ms current injections (Figure 4A, B). To determine whether fluctuating levels of estrogen are necessary to maintain post-puberty firing properties, the ovaries were removed either at the start (3 weeks: 3w OVX) or at the end (6 weeks: 6w OVX) of puberty and recordings were carried out at 7w. In both groups, estrogen implants that should provide a constant physiological level (27) were given at 6 weeks to mitigate the hyperexcitability of *Kiss1*^*ARC*^ neurons that is otherwise caused by entirely removing estrogen-mediated negative feedback.

**Figure 4.**
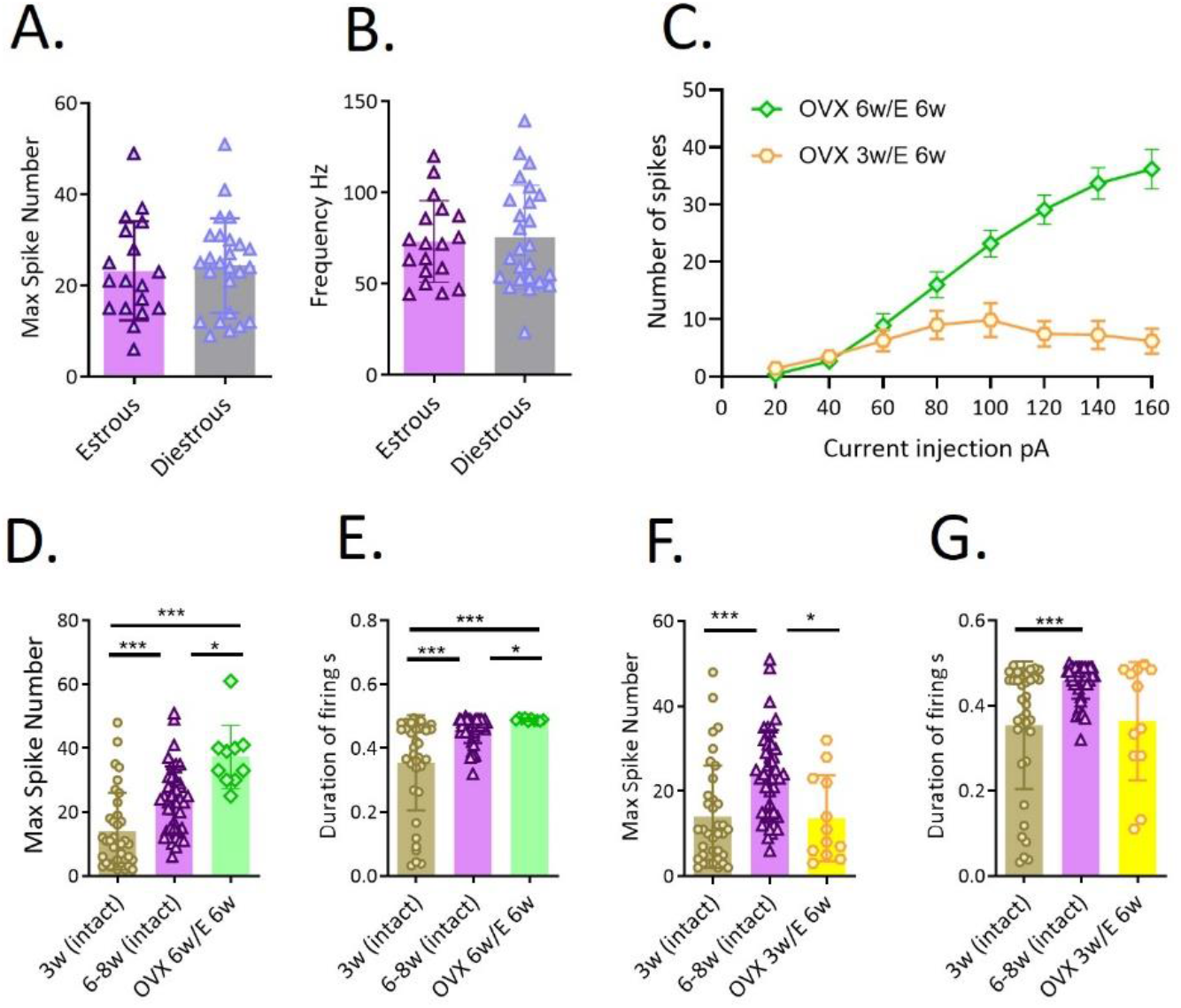
Intrinsic plasticity in *Kiss1*^ARC^ neurons is partially prevented by OVX at 3 weeks. A, B. Maximum number of spikes (P=0.7) and maximum firing frequency (Hz; P=0.86) in *Kiss1*^*ARC*^ neurq 6-8w post-puberty female mice in the estrous phase (lilac bars) or diestrous phase (grey bars) of the estrous cycle). C. The number of spikes versus current injection in *Kiss1*^*ARC*^ neurons from mice that had OVX at 3w (yellow hexagons, n = 12) or at 6w (green diamonds, n = 10); in both groups, estrogen implants were given at 6w. Mixed model: Current, P<0.0001, F(7,167)=14; Age, P<0.0001, F(l,167)=77. D, E. Maximum spike number and maximum duration of firing in mice that had OVX at 6w, compared with 3w intact and 6-8w intact mice (D and E, Kruskal Wallis, P<0.0001. F, G. Maximum spike number and Maximum duration of firing in mice that had OVX at 3w compared with intact 3w and 6-8w mice (F, Kruskal Wallis, P<0.0001; G, Kruskal Wallis, P=0.0004). All posthoc P values: *P<0.05; **P<0.01; ***P<0.001.

The number of spikes per current injection in *Kiss1*^*ARC*^ neurons from 3w OVX mice compared with 6w OVX mice is shown in Figure 4C; there was a significant effect of both age and current injection, with profoundly lower numbers of spikes in 3w OVX mice. Firing properties were compared with those in intact 3 week and intact 6–8-week-old *Kiss1*^*ARC*^ neurons. In 6w OVX mice, the maximum number of spikes was significantly larger than both 3w and 6w intact mice; this likely reflects residual hyperexcitability, despite the estrogen implant at 6w (Figure 4C). Note that 6w OVX mice do not resemble 3w intact mice, indicating that OVX at 6w does not prevent intrinsic plasticity occurring in *Kiss1*^*ARC*^ neurons. The duration of firing was significantly different in 6w OVX mice compared with intact 3w and 6-8w mice (Figure 4E) but had a similar mean value to intact 6-8w mice.

In 3w OVX mice, the maximum number of spikes recorded at 7w in *Kiss1*^*ARC*^ neurons was the same as in 3w intact mice, but significantly lower than the maximum number of spikes in 6-8w intact mice (Figure 4F), while the duration of firing was intermediate between 3w and 6-8w intact mice (Figure 4G). This suggests that OVX at 3w, with the resulting absence of estrogen at 3-6 weeks, prevents the intrinsic plasticity occurring in *Kiss1*^*ARC*^ neurons at puberty, such that *Kiss1*^*ARC*^ neurons fail to achieve their mature physiological phenotype.

## Discussion

Neuronal plasticity is associated with critical periods of brain development, and the most important developmental period for the survival of species is puberty, when individuals achieve fertility and reproductive capacity. We report for the first time evidence that hypothalamic *Kiss1*^*ARC*^ neurons, that control puberty and fertility, undergo intrinsic neuronal plasticity at puberty in female mice, including changes in action potential firing properties and changes in Na_V_, K_V_ and HCN ion channel genes. Prior to puberty, in 3-week-old mice, *Kiss1*^*ARC*^ neurons have firing properties that would make them unsuitable for the sustained activity that is needed to activate GnRH neurons and trigger LH secretion in the HPG axis. Consistent with this, very few GnRH/LH pulses are detected at weaning (3w) in mice, but this increases during the juvenile period (3 to 4w) (32). The changes we have identified could endow post-puberty *Kiss1*^*ARC*^ neurons with a mature physiological phenotype that allows them to fire in a sustained manner. This would be amenable to neuropeptide modulation for generation of burst firing that results in pulsatile release of kisspeptin from *Kiss1*^*ARC*^ neurons and that allows them to perform their role as activators of the HPG axis (33). Consistent with this, GnRH neurons stimulated optogenetically for 2 min at 10 Hz elicit LH pulses resembling physiological activity (21, 34). Intrinsic plasticity of action potential firing in *Kiss1*^*ARC*^ neurons begins between 3-4 weeks of age, just prior to puberty onset, and appears to require ovarian estrogen to be present at least between 3-6 weeks of age.

The passive and active membrane properties of *Kiss1*^*ARC*^ neurons are well characterised in adult female mice (35). They are small neurons with relatively low capacitance and high input resistance and a resting membrane potential of -50 to -75 mV (7-14) reviewed by (35, 36). Most of our recordings were carried out in neurons from intact pre- and post-puberty female mice; no differences in the capacitance and input resistance or resting membrane potential were observed, indicating that these do not change at puberty.

Many *Kiss1*^ARC^ neurons show no spontaneous action potential firing, while others show irregular firing with highly variable frequency (0.5-5 Hz) (7-14). To our knowledge, only one study describes spontaneous action potential firing of pre-pubertal *Kiss1*^ARC^ neurons (P18-25), showing that, similar to post-pubertal *Kiss1*^*ARC*^ neurons, 70% are silent while 30% fire irregularly at around 2.5 Hz (10). Even less is known about the firing of *Kiss1*^*ARC*^ neurons in response to controlled injections of depolarising current, although in OVX female mice frequencies of up to 20Hz were seen with 90-120pA of current injection in NKB neurons (12). In *Kiss1*^*ARC*^ neurons from adult male mice, firing up to 40Hz was observed along with an outward potassium current mediated by K_V_4.2; this increased firing irregularity in response to depolarizing current (14). Many electrophysiology studies of *Kiss1*^*ARC*^ neurons have been done in OVX or OVX+E female mice to capture their phenotype when there is no cycling of sex hormones. *Kiss1*^*ARC*^ neurons from OVX mice are more excitable compared to neurons from OVX+E mice or intact mice. Their action potential threshold (9, 37), duration of firing (37), firing frequency (12), and spontaneous action potential firing (8) are all significantly higher. To standardise measurements of firing between pre- and post-puberty mice, we held the membrane potential at - 75mV and delivered depolarising current injections to determine the properties of repetitive firing for 500 ms. In these recordings, *Kiss1*^*ARC*^ neurons showed a dramatic change in firing properties, from high frequency short duration and irregular firing in mice aged 3 weeks to sustained firing at lower frequency with greater regularity by 6-8 weeks.

The short duration firing at 3 weeks could be explained by depolarisation block: a period of silence in the depolarised neuron, characterised by high frequency firing that leads to inactivation of Na_V_ channels (38). Consistent with this, firing was more susceptible to a decrease in spike number with increasing current injection. There might also be spike frequency adaptation: an initial higher frequency slows down during a train of spikes, sometimes stopping altogether (39). The ratio of the last/ first inter-spike intervals was greater in 3-week-old mice when compared with 6-8 weeks. In general, there was more variability in all parameters measured in *Kiss1*^*ARC*^ neurons from 3-week-old mice, suggesting that individual neurons are developing at a different rate and some achieve a more mature phenotype earlier than others. None-the-less, the overall population was significantly different from those in 6–8-week mice, and even in 4-week mice, in a number of measures (Figure 1). The changes in firing properties could allow post-puberty *Kiss1*^*ARC*^ neurons to fire action potentials over a longer period, providing a convenient time window for modulation by NKB and dynorphin for pulsatile release of kisspeptin and thus GnRH and LH (40).

Evaluation of the action potential waveform revealed several potential explanations for the changes in repetitive firing at puberty. The threshold for action potential generation as well as the trough following the action potential were both more hyperpolarised in post-puberty mice, and these changes correlated with the increase in spike number seen in 6–8-week-old mice. It is possible that K_V_ channels contributing to action potential repolarisation and the after-hyperpolarisation (represented by the trough) might make firing frequency slower and steadier, allowing for more sustained firing. In addition, changes in the upstroke of the action potential might change the excitability of the neurons. Consistent with this, changes were seen in the expression of genes encoding Na_V_1.2 channels (*Scn2a*, which shows higher expression in the adult mouse hypothalamus than *Scn1* or *Scn3*; Allen Mouse Brain Atlas), K_V_7.2 (*Kcnq2*) channels and the γ3 but not α subunit genes of the BK channel. While a decrease in *Scn2a* alongside an increase in spike number at puberty might seem counter-intuitive, Na_V_1.2 channels have been shown to cause neurons to fire fewer spikes (41). Post-puberty *Kiss1*^AVPV^ and *Kiss1*^ARC^ neurons express the Na_V_ channel genes *Scn1a, Scn2a*, and *Scn6a* subunits in estrogen-replaced OVX mice (42), and so further changes in Na_V_ channel genes might occur at puberty.

The changes in firing in *Kiss1*^*ARC*^ neurons at puberty might also be due to changes in BKγ3 or K_V_7 potassium channels. BK channel auxiliary subunits (BKγ1, BKγ2, and BKγ3) shift the voltage dependence of activation to more hyperpolarised potentials, facilitating activation (43); this might suggest that in *Kiss1*^*ARC*^ neurons from post-puberty mice, expressing less *Lrrc55* (encoding BKγ3) is unlikely to account for a larger trough/ fast AHP. Adult female *Kiss1*^ARC^ neurons have M current mediated by K_V_7 channels, which regulates the resting membrane potential (44).

One further change at puberty was the appearance of a larger amplitude mAHP following a train of action potentials, and greater prevalence (seen in most post-puberty neurons, but in only half of pre-puberty neurons). A mAHP can also provide a pause in firing to prevent hyper-excitability and depolarisation block and assist with removal of channel inactivation. Ion channels that usually contribute to the mAHP include small conductance calcium-activated K^+^ channels and K_V_7 channels; in this study, genes encoding SK2 and SK3 were not changed after puberty. This does not mean their expression is not changing; a potential caveat is that the mAHP is detected in 50% of 3-week-old mice and 70% of 6–8-week-old mice, and as such detecting a change in gene expression is likely to be challenging. Combined inhibition of SK and BK channels significantly inhibited spike frequency adaptation in *Kiss1*^ARC^ neurons of adult male mice, so that SK and/or BK channels were postulated to limit excitability during high frequency firing (45); thus, SK channels could contribute to the mAHP. Genes encoding K_V_7.2 and K_V_7.5 were either decreased or unchanged, respectively; suggesting they do not contribute to the mAHP. HCN channels, a family of non-selective voltage-gated cation channels whose activity influences hyperpolarization time and rebound firing (46), can contribute to the depolarisation following a mAHP, and this might be attributed to HCN1 channels in *Kiss1*^*ARC*^ neurons as *Hcn1* gene expression increased at puberty. *Hcn1-4* was also detected in adult *Kiss1*^*ARC*^ neurons by single cell qPCR (7). Additionally, functional hyperpolarization-activated cyclic nucleotide-gated (HCN) channels have been demonstrated in adult mouse (47) and guinea pig (48) *Kiss1*^*ARC*^ neurons.

The changes seen in post-puberty mice could be explained by the simple presence of estrogen, which begins to rise prior to puberty onset. If changes in estrogen concentration *per se* cause the increase in spike number, decrease in frequency and sustained firing, it is likely that firing properties in response to current injection would change across the estrous cycle, and certainly between estrous, when estrogen levels are higher, and diestrous (31). Yet, there were no differences in any repetitive firing properties between estrous and diestrous mice, suggesting that peaks in estrogen concentration do not account for the firing properties in mature *Kiss1*^*ARC*^ neurons. A further possibility is that the cycling of estrogen during the estrous cycle is necessary for intrinsic plasticity to occur. To test this, we performed OVX at the end of puberty to prevent cyclical release of estrogen (and supplemented with estrogen implants to restore the negative feedback and prevent hyper-excitability of *Kiss1*^*ARC*^ neurons). OVX at 6 weeks did not prevent *Kiss1*^*ARC*^ neurons from acquiring a mature firing phenotype; firing resembled that seen in intact 6-8w mice but was higher in spike number due to compromised estrogen-mediated negative feedback. Conversely, OVX at the start of puberty (3 weeks), with no estrogen supplement until 6 weeks, led to firing properties in post-puberty mice that more closely resembled those seen in 3-week pre-puberty mice, suggesting that OVX had arrested the developmental plasticity needed to achieve a mature *Kiss1*^*ARC*^ neuronal phenotype. One possibility is that estrogen is needed during a critical period at 3-6 weeks for intrinsic plasticity to occur, but since we have not ruled out a role for progesterone we cannot claim sufficiency.

The changes in firing observed in this study occur in parallel to changes in *Kiss1* mRNA levels in female rats which show a fourfold increase at 3-4 days before vaginal opening (49). Thus, intrinsic plasticity along with changes in *Kiss1* expression might be critical changes occurring in *Kiss1*^*ARC*^ neurons as they adapt to adult reproductive functions. The two may occur independently or be inter-related; for example, sustained firing post-puberty may require more kisspeptin synthesis for release onto GnRH neurons.

In conclusion, post-pubertal firing with higher number of spikes, lower frequency and more regularity might represent the mature physiological phenotype needed for *Kiss1*^*ARC*^ neurons to act as the pulse generators for GnRH and LH release. Pre-pubertal *Kiss1*^*ARC*^ neurons, which fire with a lower number of spikes, higher frequency and more irregular, may represent an immature phenotype which cannot support *Kiss1*^*ARC*^ neurons in their reproductive functions. Changes in the expression of specific ion channels could underlie this neuronal plasticity, and depend on the presence of estrogen during a critical window of pubertal development. A similar phenomenon has been reported in parenting circuits, where sex steroid hormones during pregnancy rewire the medial preoptic area of the mouse hypothalamus (50). It will be interesting to explore whether and how intrinsic plasticity in *Kiss1*^*ARC*^ neurons is modulated by environmental factors that influence puberty onset, such as social isolation (51) and nutrition (52) to determine whether they can delay or accelerate *Kiss1* neurons intrinsic neuronal plasticity.

